# When sleep reports diverge from physiology: correlates of sleep discrepancy in a health population

**DOI:** 10.64898/2026.08.24.746676

**Authors:** Péter Przemyslaw Ujma, Róbert Bódizs

**Affiliations:** Semmelweis University, Institute of Behavioural Sciences

**Keywords:** sleep discrepancy, convergent validity, personality, insomnia, multiday observational study

## Abstract

**Study Objectives:** Sleep characteristics are often estimated using self-reports, which may differ from objective measurements, a phenomenon known as sleep discrepancy. However, the factors associated with the degree of sleep discrepancy remain poorly understood.

**Methods:** In the current study, a large healthy participant sample of the Budapest Sleep, Experiences and Traits Study (total N=267, 1899 nights) completed a 7-day protocol including mobile EEG recordings and sleep diaries, and also provided questionnaire-based reports of habitual sleep. We compared analogous sleep metrics from these three modalities, and used cross-validated LASSO regression to investigate demographic, psychological and lifestyle-related factors associated with increased sleep discrepancy across all three modalities, at both between- and within-participant levels.

**Results:** Daily diaries estimated EEG-based sleep timing accurately (mean r=0.83), but were less accurate for sleep onset latency and quality. In contrast, questionnaire reports of habitual sleep provided inaccurate measures of even sleep timing (mean r=0.49) and considerably misestimated sleep timing and duration. Insomnia and depressive symptoms, napping, co-sleeping and personality traits were associated with increased sleep discrepancy.

**Conclusion:** In healthy adults, questionnaires about habitual sleep provide only moderately accurate and biased estimates of actual sleep. Daily diaries provide considerably more accurate estimates, but sleep onset latency and physiological sleep quality is estimated by all self-reports less accurately than sleep timing. Sleep discrepancy is also present in healthy participants, it is particularly and its degree is affected by non-pathological characteristics. Long-term monitoring by daily diaries or wearables should be preferred to self-report questionnaires to measure sleep.

**Statement of significance:** This study provides a comprehensive evaluation of how closely two types of subjective reports (habitual sleep questionnaires and daily sleep diaries) track EEG-based objective sleep in a large community sample. We show that while diaries estimate sleep timing accurately, questionnaires about habitual sleep are both inaccurate and biased. The degree of objective-subjective sleep discrepancy varies both across and within-individuals as a function of depressive and insomnia symptoms, sleep habits, and personality traits. The findings show that self-reports of habitual sleep provide inaccurate and distorted estimates of sleep behavior, especially in participants with subclinical psychiatric symptoms. Longitudinal diary- or wearable-based assessments are increasingly viable and more valid alternatives.

## Introduction

Sleep problems are common, constitute a substantial epidemiological and economic burden ^1^, and are frequently undiagnosed ^2,3^. In both clinical and community settings, sleep is measured using various methods with a general tradeoff between simplicity and accuracy.

The conceptually simplest questionnaires require participants to report their habitual sleep quality ^4^ or sleep timing ^5,6^. Sleep diaries constitute another method and require repeated self-reports of the quality and timing of recent sleep ^7,8^. Actigraphy and other wearables ^9–11^ especially mobile EEG systems ^12,13^ and polysomnography ^14^ objectively measure the timing and physiological quality of sleep, including the staging of sleep which is normally not accessible to self-reports. Instrument-based sleep measures, however, do not capture the subjective experience of sleep, which is imperfectly related to physiological sleep quality ^15–17^, with an independent epidemiological significance ^18,19^.

While there is a considerable correlation between self-reported and objectively measured sleep metrics ^16,20–23^ sleep measures by various methods are in imperfect agreement, a phenomenon referred to as sleep discrepancy ^24^, sleep misperception ^25^, or, in the case of clinically significant subjective sleep complaints which are not verifiable by objective measures, paradoxical ^26^, subjective ^27^, or pseudo-insomnia ^28^. Discrepancies between subjectively and objectively assessed sleep are especially common in sleep disorders. In insomnia, sleep duration and latency are frequently overestimated while sleep efficiency is underestimated by self-reports compared to objective measures ^29–32^. A similar underestimation of sleep quality is associated with depressive symptoms ^33,34^. In contrast, in obstructive sleep apnea, patients are often unaware of and underreport the sleep fragmentation characteristic for this illness ^35,36^. Other characteristics predicting sleep discrepancy were less systematically researched, but include psychosocial characteristics ^37^, general health and functioning in the elderly ^31^, personality ^38^, nightmares in PTSD ^39^ and various psychiatric symptom profiles ^40^.

Most research on the causes of sleep discrepancy focuses on between-participant effects, that is, the discrepancy between average self-reported and objectively measured sleep. Some research is, however, available on the within-participant correlates of sleep discrepancy, that is, factors associated with the same person providing self-reports more or less congruent with objective measures on different days. Herbert et al ^41^ in a small sample of participants with insomnia symptoms found that high pre-sleep cognitive activity, sleep effort and low morning mood predicted a greater underestimation of sleep time and overestimation of sleep onset latency. Acute symptom severity also predicted poor subjective reports of sleep in fibromyalgia patients ^42,43^, while in a healthy sample psychological attitudes predicted discrepancies ^44^.

Despite the high frequency of sleep complaints in the general population in the absence of a formal diagnosis ^2,3^, most research about sleep discrepancy is performed in clinical samples, and investigates a limited, symptom-focused list of potential predictors of the mismatch between subjectively reported and objectively measured sleep. In the current study, we leveraged a large sample of healthy volunteers with a seven-day observation period incorporating mobile EEG to estimate how accurately subjective sleep reports track analogous objective metrics, and a large pool of potential predictors used in cross-validated models to investigate what trait-like characteristics or daily experiences account for their discrepancy.

## Methods

### Data

Data from the Budapest Sleep, Experiences and Traits Study (BSETS) was used. The full BSETS protocol is available in a separate publication ^45^. Briefly, BSETS is a multiday observational study in which healthy volunteers slept with mobile EEG devices on at least seven consecutive days, recorded their daily experiences each evening and subjective assessment of nightly sleep each morning via diaries. Participants were recruited using branched diffusive convenience sampling, mostly from students of Semmelweis University and relatives. The mean age was 28.93 years (SD=12.77 years, range: 18-76 years). 45% of participants were male and 55% female. In total, 267 participants completed at least some part of the BSETS protocol on 1899 days/nights. Minor deviations in these figures from previous BSETS papers reflect the discovery of new data and corrections. More precise valid Ns accounting for missingness in specific variables (typically <10%) are reported in the Results.

As maximal ecological validity was a key goal in designing BSETS, inclusion criteria were minimal and consisted only of non-recruitment from clinical settings, no intervention was given and participants were encouraged to proceed with the normal course of their lives without any restrictions on waking or sleep behavior.

An extensive demographic and psychological test battery was administered on a separate occasion to assess trait-level characteristics.

The Institutional Review Board (IRB) of Semmelweis University, as well as the Hungarian Medical Council (under 7040-7/2021/ EÜIG “Vonások és napi események hatása az alvási EEG-re” [The effect of traits and daily activities and experiences on the sleep EEG]), approved BSETS as compliant with the latest revision of the Declaration of Helsinki. All participants gave written informed consent on a form reviewed and approved by the IRB.

### Sleep measures

Throughout the study, we analyze the accuracy (correlations) and bias (mean differences) of analogous sleep metrics obtained by habitual sleep reports, daily diaries, and objective mobile EEG measures.

Habitual sleep was assessed as part of the demographic and psychological test battery, temporally separate from the participants seven-day sleep measurement protocol. The timing of habitual sleep was obtained from the Hungarian version ^46^ of the Munich Chronotype Questionnaire (MCTQ) ^6^. The quality of habitual sleep was assessed by the Hungarian version ^47^ of the Athens Insomnia Scale ^48^.

Custom diaries ^45^ were administered each evening and each morning during the seven-day sleep protocol. In evening diaries, participants recorded experiences during the previous day which were used as predictors of sleep discrepancy (see Results for details) In the diaries, participants reported the time they went to sleep on the previous night, the time (in minutes) it took them to fall asleep, and their rise times the next morning. Sleep quality was assessed using the Hungarian version ^49^ of the Groningen Sleep Quality Scale (GSQS). In morning diaries, participants also recorded co-sleeping, sleeping at strange places, and recalling dreams, which were used as predictors of sleep discrepancy.

Sleep EEG recordings were performed during each night of the sleep protocol with the Dreem2 headband. Dreem2 uses dry silicone electrodes at the positions Fp1, F7, F8, O1 and O2 (plus a bias electrode) to record EEG with a sampling frequency of 250 Hz ^45,50^. Clock time at recording start/end is logged, and a validated ^51^ complimentary algorithm uses the resulting signal to score sleep. Algorithm estimates of sleep onset latency, wake after sleep onset, total sleep time, sleep efficiency and clock time at the last sleep epoch were compared to subjective reports, as reported in the following section.

### Sleep variables

The following sleep metrics were comparable between the three measurement methods: 1) bedtime, 2) sleep onset latency, 3) sleep onset time, 4) rise time, 5) total time in bed, 6) total sleep time, 7) chronotype, 8) social jetlag and 9) sleep quality. The operationalization of these variables is described below.

Bedtime in EEG recordings was considered as the start of the recording (“lights off”). In diaries and in the MCTQ, bedtimes were explicitly recorded.

Sleep onset latency in EEG recordings was calculated as the time (in minutes) from the start of the recording to the first non-wake epoch. In diaries and the MCTQ, this was self-reported. Due to the skewed distribution of this variable ^52^, all measures of sleep onset latency were log-transformed.

Sleep onset time in EEG recordings was the time at the start of the first non-sleep epoch. In diaries and the MCTQ, this was calculated by adding sleep onset latency to bedtime.

Rise time in EEG recordings was the end of the recording. In diaries and MCTQ, it was self-reported explicitly.

Total bedtime in EEG recordings was calculated as the mean recording duration. In diaries, it was calculated as the total time from bedtime to rise time. In the MCTQ, total bedtime was calculated as the total time from bedtime to getting out of bed.

Total sleep time in EEG recordings was objectively measured as the total time from the first to last sleep epochs. In order to maximize similarity to self-reports, wake after sleep onset (WASO, 21.01 minutes on average across all recordings) was added to the usual EEG-based TST value. In diaries, total sleep time was calculated as time from bedtime to rise time minus sleep onset latency, and in the MCTQ as time from bedtime from getting out of bed minus sleep onset latency and sleep inertia.

Chronotype from EEG recordings and diaries was calculated based on MCTQ formulas ^6^ using midsleep values. If sleep duration on weekends was no longer than on weekdays, the midpoint of the first and last sleep epochs (EEG) or the midpoint of sleep onset time and rise time (diary) was used as the estimate of chronotype. If sleep duration was longer on free days, this was corrected for oversleeping by subtracting half the difference between free day and weekly average sleep durations. In the MCTQ, chronotype was calculated using the usual formula based on working day and free day midsleep ^6^. Participants reporting shift work or waking up with an alarm clock even on free days in the MCTQ were excluded, including from EEG and diary-based analyses.

Social jetlag was calculated as the absolute difference between free day and working day midsleep.

Sleep quality was estimated with sleep efficiency from EEG data, the total score of the Groningen Sleep Quality Scale in morning diaries and the total score of the Athens Insomnia Scale in trait data. Because sleep efficiency is a skewed variable, it was reflected and log transformed (SE_transf_=log10(100-SE)). On this reversed scale, higher values of sleep efficiency reflect poorer sleep quality, as in GSQS and AIS scores.

### Statistics

Statistical analysis consisted of three steps: participant-level analyses, day-level analyses, and discrepancy analysis.

In participant-level analyses, we calculated correlations (accuracy or convergent validity) between self-reported habitual sleep metrics in the MCTQ and AIS, and analogous objectively measured (via EEG) or self-reported (via diaries) sleep metrics based on daily observations. To keep daily observation-based sleep metrics comparable to the MCTQ, they were averaged across all weekdays (and compared to working day MCTQ data) and weekends (and compared to free-day MCTQ data) recorded in BSETS. This way, six values (EEG, diary, and MCTQ on weekdays on weekends) of each sleep metric were obtained.

Chronotype and social jetlag are not defined separately for weekdays and weekends, and trait-like sleep quality was measured with the AIS which, unlike the MCTQ, does not provide separate working and free day estimates. Therefore, for chronotype, social jetlag and sleep quality we did not run separate weekday and weekend analyses. Pearson correlations between all combinations were calculated to estimate accuracy/convergent validity.

Additionally, paired-sample t-tests were calculated to estimate systematic differences in the mean values (bias). Because self-reports of sleep quality are on a subjective scale, means cannot be directly compared. Therefore, we z-transformed sleep quality variables to constrain mean differences to 0 while still allowing correlational analyses ^53^.

In day-level analyses, an analogous strategy was followed, but sleep metrics were centered within participants before accuracy analyses, capturing the degree to which objective and subjective reports of sleep covary within the same person. Because due to this centering mean differences were uninformative, only correlations were calculated. As the MCTQ only provides participant-level data, day-level analyses only used EEG- and diary-based sleep metrics.

Discrepancy analysis aimed to uncover which participant-level or day-level variables are associated with the discrepancy between objectively measured and self-reported sleep metrics. A comprehensive list of participant-level (trait-like, for participant-level analyses) or day-level (time-variant, for day-level analyses) BSETS variables was used as predictors in LASSO regression ^45,54^, with the absolute difference between each EEG-based and self-rated sleep metric as the dependent variable. LASSO is a simple learning algorithm which uses increasing penalties to non-zero regression weights to select a linear combination of N variables from K>>N total variables which best predict an outcome in a holdout sample ^55,56^. LASSO was implemented with the lasso() MATLAB function, using five-fold cross-validation. The model with the lowest mean squared error was selected as optimal. In order to increase the reliability of findings, LASSO was repeated 1000 times for each metric, only predictors which were selected in the optimal model in at least 80% of runs were retained, and their coefficients were set to be the average across all runs in which they were selected. Additionally, for day-level analyses where a larger number of observations was available, a separate validation sample comprising 20% of all valid observations was randomly selected and used to test the predictive accuracy of the optimal model. This way, the use of LASSO enabled a hypothesis-free, cross-validated exploration of what individual or day-level characteristics are associated with a larger absolute discrepancy between objectively measured and self-reported sleep.

In participant-level discrepancy analyses we used the following standard ^45,54^ list of trait-level variables recorded in BSETS (measurement instruments in parentheses): age, sex, body mass index (self-reported), positive and negative emotionality (7-day averages of PANAS subscales), intelligence (ICAR-16), depression (PHQ-9), alexithymia (TAS), insomnia symptoms (AIS), dissociation (DES), the personality traits sociability, activity, aggression, neuroticism, and impulsive sensation seeking (ZKPQ), extraversion, agreeableness, conscientiousness, neuroticism and openness (BFI-44), and 7-day average nap frequency, nap duration, co-sleeping frequency, and frequency of sleeping at unfamiliar places. Both predictors and differences were expressed as z-scores to make regression coefficients directly comparable and interpretable as correlation coefficients. Because the quality of habitual sleep was operationalized by AIS score, this variable was not included as the predictor of discrepancies.

In day-level discrepancy analyses, EEG-diary differences of analogous sleep metrics were calculated first. Then, the differences were centered within each participant, and the centered differences were z-transformed. This process ensures that only within-participant differences inform analyses (due to centering) but ensures comparable effect sizes across sleep metrics and preserves between-participant differences in the raw discrepancy (due to the z-transformation of pooled differences). The following day-level variables were used as predictors (after similar z-transformations) of within-person EEG-diary differences: weekend night, co-sleeping, sleeping in a foreign place, reporting a dream, Groningen Sleep Quality Scale total score, a checklist of 13 self-reported daily experiences during the day preceding the night ^45^, 10 positive and negative previous-day emotions reported using the short-form PANAS ^57^, and Likert scale ratings of the previous day as happy, interesting, mentally and physically exhausting ^45^. Note that because the MCTQ only provides participant-level data, day-level analyses only used EEG and diary reports. The large number of day-level associations enabled the use of an independent validation sample (20% of all observations). Models were retained if the degree of freedom-adjusted out-of-sample R^2^ of the optimal model was >0, that is, if the model predicted meaningful additional variance.

In order to control for multiple comparisons, in bias analyses (mean differences of analogous sleep metrics), a Benjamini-Hochberg correction for false discovery rates ^58^ was applied to the entire pool of p-values obtained from paired-sample t-tests (Table 1). In accuracy analyses (correlations of analogous sleep metrics), statistical significance was of secondary importance as a mere rejection of the null hypothesis of zero correlation is insufficient to support convergent validity, which requires a high effect size. In these analyses, all correlations are highly significant (p<0.001) unless otherwise indicated.

**Table 1.**
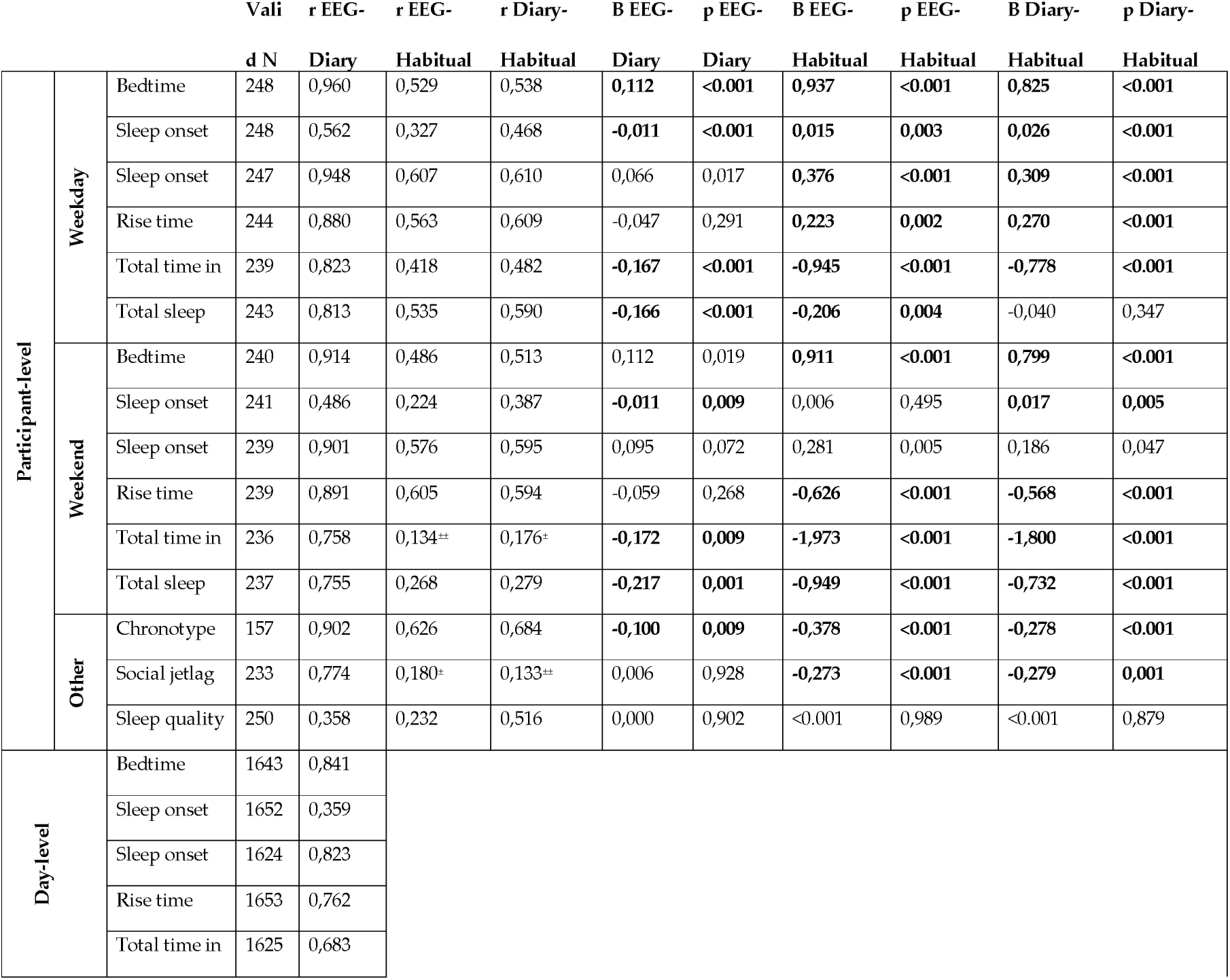
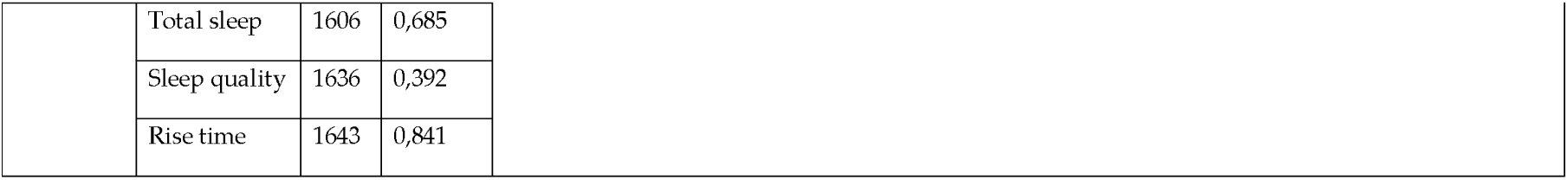
Detailed statistics about accuracy and bias. Participant-level (trait-level correlates of the mean discrepancy of participants) and day-level (day-level correlates of within-person differences in discrepancy) discrepancy is reported separately. Weekday and weekend estimates are provided separately for participant-level analyses which use the MCTQ. Valid N indicates the number of observations with full data. Correlations (r) indicate the similarity of analogous sleep metrics. Regression coefficients (B) indicate the discrepancy of the means, expressed in logged minutes for sleep onset latency and hours for all other metrics. Positive values indicate higher means in the first-listed data source. Bolded values indicate significant mean differences after FDR correction. All correlations are p<0.001 and significant after FDR correction unless otherwise indicated: ^±^p<0.01, ^±±^p<0.05.

## Results

### Participant-level analyses

Diaries estimated sleep timing accurately with little bias, but were less accurate about sleep onset latency and sleep quality. Habitual sleep reports were less accurate even about timing and exhibited greater bias.

Specifically, on weekdays, EEG- and diary-based measurements showed high agreement (mean r=0.83). The lowest agreement was on sleep onset latency (r=0.56), and the highest on bedtime (r=0.96). The agreement with habitual sleep reported in the MCTQ was moderate (mean r_EEG_=0.53, mean r_diary_=0.54). Despite the high agreement with EEG, diaries underestimated bedtime by ∼6.6 minutes, overestimated sleep onset latency by ∼1 minute and total sleep time by ∼10 minutes. MCTQ-based bedtimes were close to a full hour and sleep onset times over 20 minutes later than either diary- or EEG-based measures.

Participants also reported earlier habitual rise time in the MCTQ than what was recorded with EEG or reported diaries, but because the difference was smaller at <20 minutes, total sleep time and especially total bedtime were also substantially higher in the MCTQ than in daily recordings. Figure 1 illustrates the similarity of weekday sleep metrics across the three measurement modalities.

**Figure 1.**
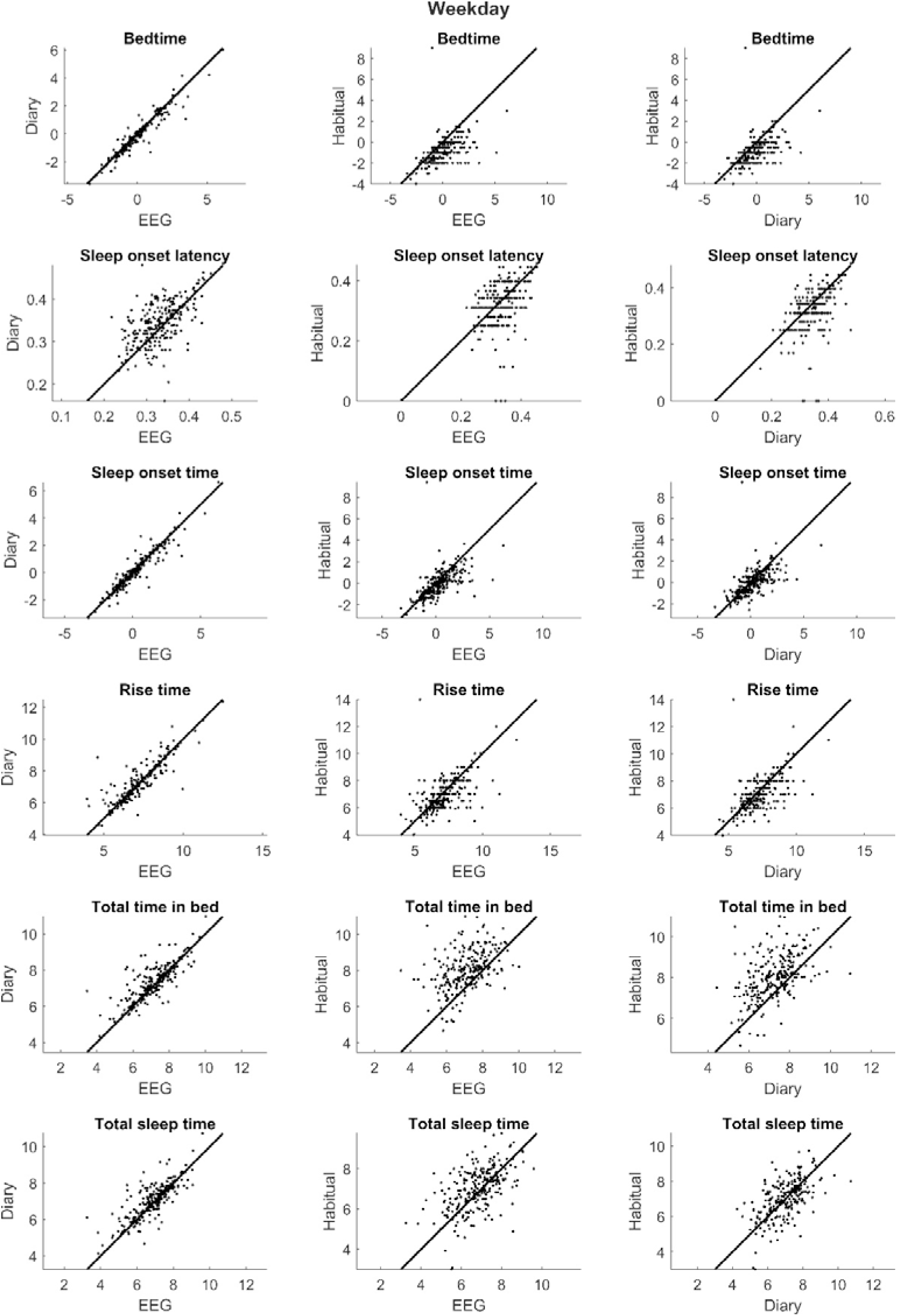
Participant-level correlations and mean differences in analogous sleep metrics on weekdays/working days, across three modalities: multiday averages of EEG measurements (EEG), multiday averages of morning diaries (Diary) and habitual sleep reported in the MCTQ (Trait). Scatterplots illustrate correlations. An identity line is shown (x=y) to highlight differences in means. Units are in hours for durations and hours relative to midnight (0) for timing.

On weekends, the agreement of various sleep metrics was nominally lower (mean r_EEG-diary_=0.78, _rEEG-MCTQ_=0.38, r_diary-MCTQ_=0.42). As during weekdays, the correlation of diary- and EEG-based sleep metrics was strongest for sleep timing and lowest for sleep onset latency and total time in bed. Compared to EEG, diaries systematically overestimated sleep time and especially time in bed (see the Limitations, however, about the poor ability of EEG to track sleep inertia which may bias these findings). As during working days, participants reported earlier free-day bedtimes in the MCTQ than those recorded with diaries and EEG, while rise times were reported to be later than actually measured, resulting in about two hours less total bedtime and close to a full hour less sleep time in diaries and EEG than in the MCTQ. Figure 2 reports the similarity of weekend sleep metrics across the three measurement modalities.

**Figure 2.**
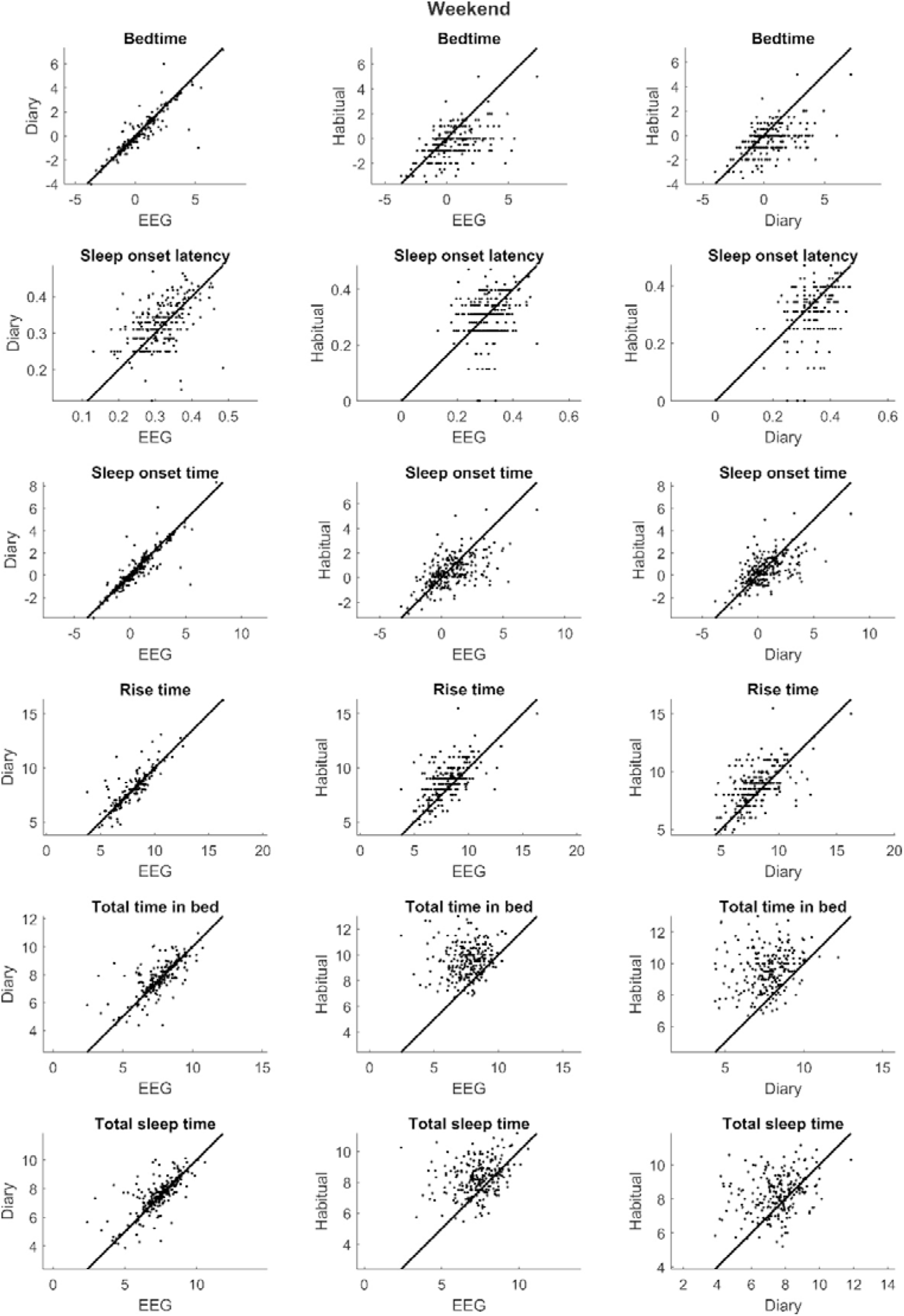
Participant-level correlations and mean differences in analogous sleep metrics on weekends/free days, across three modalities: multiday averages of EEG measurements (EEG), multiday averages of morning diaries (Diary) and habitual sleep reported in the MCTQ (Trait). Scatterplots illustrate correlations. An identity line is shown (x=y) to highlight differences in means. Units are in hours for durations and hours relative to midnight (0) for timing.

Chronotype (r=0.9) and social jetlag (r=0.77) were highly similar across EEG- and diary-based measurements. The agreement with MCTQ-based chronotype was moderate (r=0.63-0.68), but with social jetlag it was low (r=0.13-0.18). MCTQ-based chronotype and social jetlag estimates were systematically higher than those resulting from either diaries or EEG measures.

Sleep quality based on the AIS and morning GSQS scores was moderately correlated (r=0.52), while a somewhat lower correlations were observed with EEG-based sleep efficiency (r=0.23-0.36). Figure 3 illustrates the similarity of chronotype, social jetlag, and sleep quality across the three measurement modalities.

**Figure 3.**
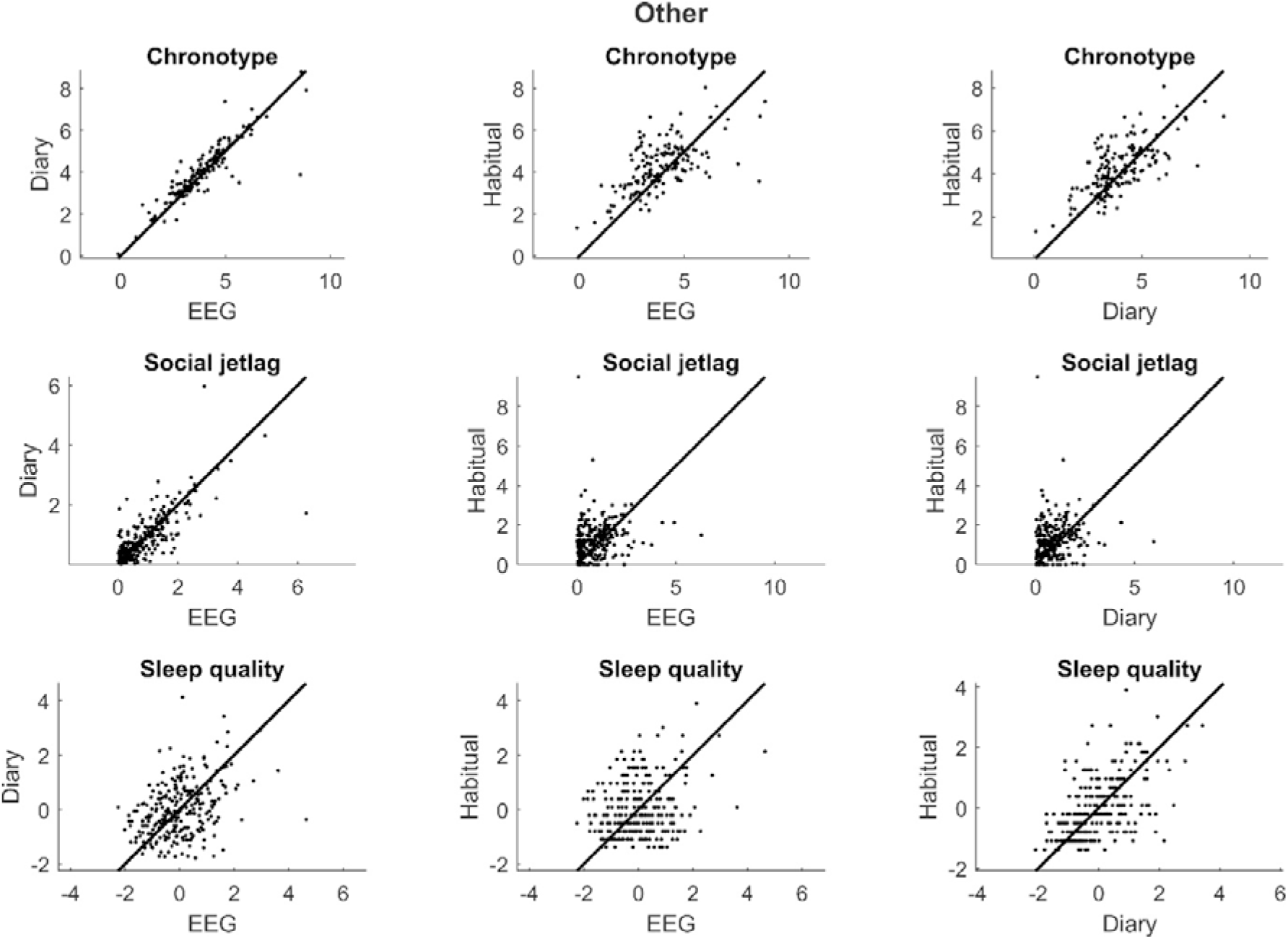
Weekend participant-level correlations and mean differences in analogous sleep metrics, across three modalities: multiday averages of EEG measurements (EEG), multiday averages of morning diaries (Diary) and habitual sleep reported in the MCTQ or AIS (Trait). Scatterplots illustrate correlations. An identity line is shown (x=y) to highlight differences in means. As sleep quality has no natural unit, values are z-scores and the mean difference is thus constrained to 0. Other units are in hours for durations and hours relative to midnight (0) for timing.

Table 1 provides detailed statistics about valid Ns, accuracy (correlations) and bias (mean differences) of sleep metrics across the three modalities.

### Day-level analyses

Day-level analyses aimed to capture how similarly the within-participant fluctuations of sleep metrics are captured by EEG and self-report diaries.

Bedtime, sleep onset time and rise time exhibited high but less than perfect correlations (r=0.76-0.84). Correlations were lower (r=0.68) for total time in bed and total sleep time, and the lowest for sleep onset latency and sleep quality (r=0.39). The findings are illustrated on Figure 4.

**Figure 4.**
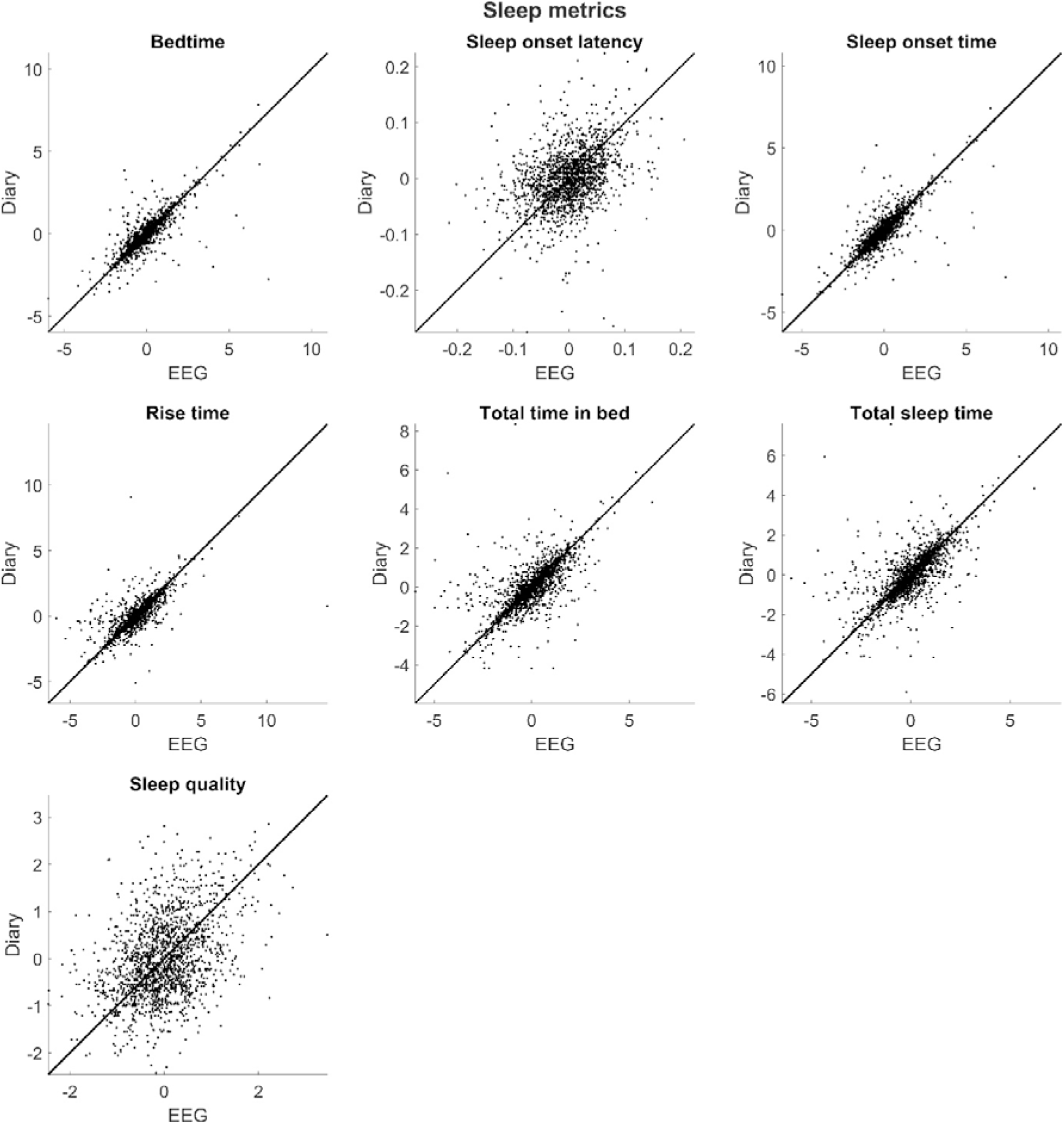
Day-level correlations and mean differences in analogous sleep metrics, across EEG and morning diaries. Data points illustrate within-participant deviations from the mean, with multiple data points per participant. Scatterplots illustrate correlations. An identity line is shown (x=y) to highlight differences in means, note, however, that in day-level analyses these mean differences are constrained to 0. As sleep quality has not natural unit, values are z-scores and the mean difference is thus constrained to 0. Other units are in hours for durations and hours relative to midnight (0) for timing.

Table 1 also reports detailed statistics about day-level sleep metrics.

### Discrepancy analysis

In discrepancy analyses, we used hypothesis-free, cross-validated LASSO regression to identify traits (in participant-level analyses) or day-variant variables (in day-level analyses) which are associated with a greater than average discrepancy between objectively measured and self-reported sleep metrics.

In participant-level analyses, a relatively large number of independent associations with sleep discrepancy were found. In habitual sleep reports compared to either EEG or diaries, participants with more insomnia symptoms overreported sleep onset latency and underreported total sleep time and time in bed (|B|=0.07-0.20). Those with depressive symptoms underestimated sleep quality (B=−0.27). Frequent nappers and those reporting more negative emotions overreported time in bed and total sleep time (|B|=0.02-0.22). Co-sleeping was associated with an underestimation of sleep onset latency (|B|=0.10). While most trait-level characteristics correlated specifically with EEG-habitual or diary-habitual sleep discrepancy, depressive symptoms led to an underestimation of sleep quality compared to EEG both in next-morning diaries (B=−0.07) and reports of habitual sleep (B=−0.27). Additionally, those with higher levels of the personality trait Openness reported less time in bed and total sleep time in either diaries or the MCTQ than recorded by EEG (B=−0.04 – −0.2). A full list of associations is provided on Figure 5.

**Figure 5.**
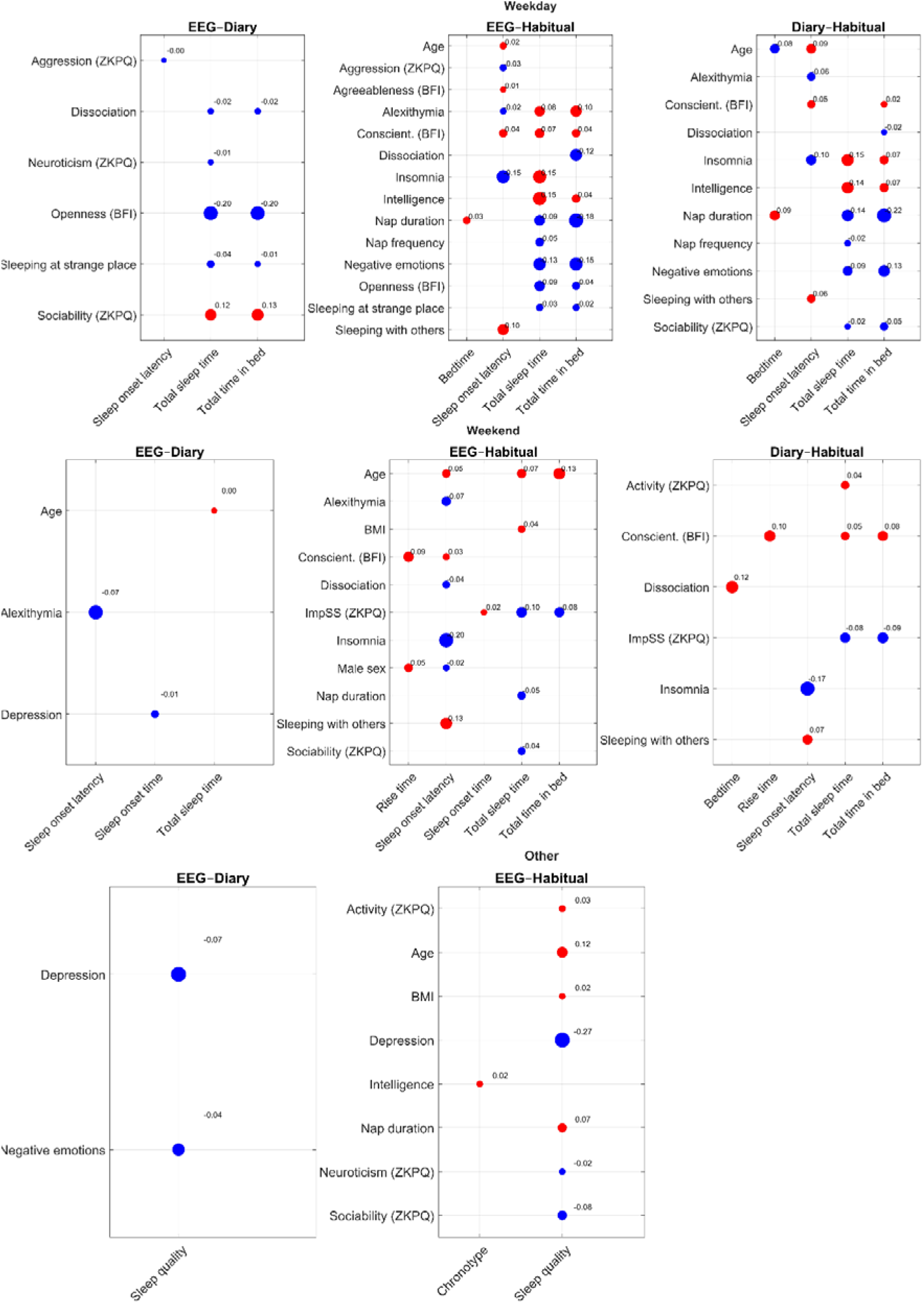
Predictors of sleep report discrepancies on weekdays/working days (top panels), weekends/free days (middle panels) and chronotype, social jetlag and sleep quality (bottom panels). Each subpanel displays the trait-level characteristics (vertical axis) predicting discrepancies of specific sleep metrics (horizontal axis) between EEG, diaries, and the MCTQ/AIS (separate panels). Dots indicate that the trait emerged as a predictor of discrepancies in the sleep metric in cross-validated LASSO models. The color of the dots indicates the sign and their size the strength of the association. A negative correlation means that having the trait characteristics displayed on the vertical axis is associated with overestimation of the sleep parameter on the horizontal axis in more subjective reports (diary/habitual>EEG or habitual>diary). The displayed regression coefficients can be interpreted as correlation coefficients as all variables are z-transformed. Predictors and sleep metrics are only shown if they were retained in at least one LASSO model. Note that sleep quality is negatively coded by our metrics so a negative correlation reflects an underestimation of sleep quality in self-reports.

In day-level analyses, only three models additional variance in the independent validation sample. Sleep onset latency (out-of-sample R^2^=0.019) was overestimated in next-morning diaries if subjective sleep quality (GSQS score) was lower in the same diaries (B=−0.05).

Current-night co-sleeping had a small independent effect (B=0.01) on underestimating sleep onset latency. Sleep quality (out-of-sample R^2^=0.01) was rated as higher if sleeping in foreign place (B=−0.025) and after public transport use (B=−0.018). (One model [total sleep time] with trivial variance accounted for [R^2^<0.019] and multiple variables retained by models with trivial effect sizes [B<0.001] are not listed but available from the data accompanying the paper).

## Discussion

In our study, we used a large sample of healthy participants undergoing a week-long sleep monitoring protocol to estimate the convergent validity of questionnaire-, diary-, and EEG-based sleep metrics, and investigate the factors associated with their discrepancy.

Sleep timing based on diaries and EEG was highly similar. Accuracy (correlations) were high, and highly EEG-congruent estimates of chronotype and social jetlag were obtained by diaries. Sleep onset latency, however, was estimated less accurately and with a slight upward bias in diaries. While diaries also underestimated bedtime and overestimated total time in bed and total sleep time, the difference was relatively small and may be partially accounted for by differences in variable definitions. This is because EEG sleep timing was based on the clock time at switching devices on and off which may not account for some time in bed recorded in diaries. The correlation between diary- and EEG-based sleep quality was moderate, in line with previous research ^15,16^, highlighting that the subjective experience of sleep depends on factors other than architecturally defined physiological sleep quality. Day-level correlations of EEG and diary data were similar to trait-level correlations. In general, these findings underscore that sleep timing is accurately recovered by next-morning diaries, although the reason sleep is perceived as restorative is imperfectly reflected by macrostructurally derived physiological sleep quality.

A different picture, however, emerged based on questionnaires of habitual sleep. In this case, accuracy was moderate at best, lower still for sleep onset latency, and high bias was present. Specifically, both during weekdays and weekends participants went to bed later and woke up earlier than the habitual sleep patterns they reported in the MCTQ, a discrepancy that persisted in comparison with either EEG or diaries. This resulted in a considerable overestimation of total time in bed and total bedtime in the MCTQ. Because bias was greater for bedtimes than for rise times, the net effect on MCTQ-assessed chronotype was positive: the chronotype based on habitual sleep was later than the chronotype estimated from EEG/diaries. While some bias may result from differences in variable definitions, this is unlikely to be a complete explanation due to its magnitude and its presence even in variables with small differences in their definition (e.g. sleep onset time which is defined analogously across all three metrics). It is also unlikely that the one-week protocol was poorly representative of participants’ habitual sleep, as most participants were students and both the protocol and MCTQ were administered during the highly structured university semester. Furthermore, participants were explicitly instructed not to modify their sleep habits. A more likely explanation is that participants had relatively little insight into the precise timing of their habitual sleep (unlike in next-morning diaries, where sleep timing was still accurately recovered from recent experience). Interestingly, diary-based sleep quality had a nominally higher correlation with habitual sleep quality (AIS scores) than with EEG-based sleep efficiency recorded during the immediately preceding night, highlighting the influence on trait-level variables (e.g. well-being, personality) on subjective reports of sleep quality. In sum, the findings indicate that self-reports of habitual sleep are relatively poorly representative of field recordings.

In an analytical step rarely taken in the literature, we constructed hypothesis-free, cross-validated multivariate models to account for the discrepancy between objectively and subjectively measured sleep. In line with the previous literature ^59–61^, we found that insomnia symptoms led to the overestimation of sleep onset latency and the underestimation of time in bed/time asleep in reports of habitual sleep. Depressive symptoms, in turn, were associated with reduced sleep quality compared to EEG in both diaries and AIS scores. These results illustrate the considerable subjective element of sleep complaints and the inadequacy of standard polysomnography-based assessment of sleep quality ^62^. One possible explanation of these findings is that negative cognitive biases are present in these disorders, which affect sleep reports even in the absence of other alterations. However, another possibility is that the relatively crude sleep architectural metrics normally available from polysomnography or mobile EEG do not fully capture the sleep alterations subjective reports reflect, and more fine-grained analyses would be necessary ^63^. Although our findings cohere with recent evidence suggesting the significant discrepancy between subjective and objective measures of sleep in insomnia patients ^61^, results on depression scores affecting sleep quality reports of the participants are less evident in the literature implying further studies and methodological considerations.

Sleeping habits emerged as another predictor of sleep discrepancy. Frequent nappers overestimated their time in bed and sleep duration, likely because they failed to account for reductions of sleep pressure due to napping. Frequent co-sleepers, in turn, underestimated sleep onset latency, possibly because they overestimated the degree to which co-sleeping synchronizes sleep.

Personality emerged as a further predictor of sleep discrepancy. The most prominent association, replicated on both weekdays and weekends and in both EEG-habitual and diary-habitual comparisons, was the underestimation of rise time and time in bed/time asleep in those with high Conscientiousness. As sleeping little and waking up early is frequently seen as virtuous, those with high values on this trait may misreport their values, with implications to research about chronotype and personality ^64,65^.

On the other hand, in day-level analyses we found limited evidence for day-variant variables predicting fluctuations of sleep discrepancy within the same person. When participants reported worse than average sleep in the morning GSQS, they overestimated sleep onset latency and sleep onset time. This is in line with the previous literature ^41^, but may contain recall bias as sleep quality and sleep timing was estimated simultaneously. No previous-day experience resulting in within-person sleep discrepancy were reliably identified.

Our work has a number of limitations. The most important limitation concerns variable definitions, which were only approximately identical across the three compared methods (questionnaires, diaries, and EEG). On the one hand, the MCTQ records sleep timing on “working days”, and “weekdays”, which we compared to EEGs and diaries recorded on weekdays and weekends, respectively. As the majority of BSETS participants were students, and 75% of participants reported exactly two free days (with another 10% three), this is likely an acceptable approximation. On the other hand, only the MCTQ explicitly asked about bedtime (separate from time at preparing for sleep) and the time of getting out of bed (separate from waking up, including sleep inertia), which may affect values of bedtime, sleep onset latency, and total time in bed, and rise time. However, very high correlations between MCTQ bedtimes and sleep preparation times (r=0.78) as well as sleep end times and times at getting out of bed (r=0.97) make it unlikely that this is a serious issue for correlations (but may impact estimates of means). A general limitation to all BSETS studies is that participants were relatively young and healthy, which strengthens the ecological validity of our findings in healthy populations but these may not generalize to clinical samples or the elderly.

## Conclusion

Our findings support the validity of diaries to estimate sleep (especially sleep timing) but reveal lower validity and higher bias in questionnaires of habitual sleep (especially for sleep onset latency). While similar tendencies were previously in clinical samples, we replicated these in a healthy population and found that the degree of sleep discrepancy is moderated by both healthy and pathological variation. Because wearable sleep measurement devices are increasingly accessible and cost-effective, their use is recommended to reduce the biases inherent in subjective reports.

## Data availability

Data and code to reproduce analyses is available at https://osf.io/j87gx.

